# Chromatin Bridges, not Micronuclei, Activate cGAS after Drug-induced Mitotic Errors in Human Cells

**DOI:** 10.1101/2021.02.02.429360

**Authors:** Patrick J. Flynn, Peter D. Koch, Timothy J. Mitchison

**Affiliations:** Department of Systems Biology, Harvard Medical School, Boston, MA 02115, USA; Center for Systems Biology, Massachusetts General Hospital, Boston, MA 02114, USA

**Keywords:** Taxanes, Anti-mitotics, cGAS, Interferon, Micronuclei, Chromatin Bridges, Aneuploidy, Inflammation

## Abstract

Mitotic errors can activate cGAS and induce type-I interferon (IFN) signaling. Current models propose that chromosome segregation errors generate micronuclei whose rupture activates cGAS. We used a panel of anti-mitotic drugs to perturb mitosis in fibroblasts and measured abnormal nuclear morphologies, cGAS localization and IFN signaling in the subsequent interphase. Micronuclei consistently recruited cGAS without activating it. Instead, IFN signaling correlated with formation of cGAS-coated chromatin bridges that were selectively generated by microtubule stabilizers and MPS1 inhibitors. cGAS activation by chromatin bridges was suppressed by drugs that prevented cytokinesis. We confirmed cGAS activation by chromatin bridges in cancer lines that are unable to secrete IFN by measuring paracrine transfer of 2′3-cGAMP to fibroblasts. We propose that cGAS is selectively activated by self-chromatin when it is stretched in chromatin bridges. Immunosurveillance of cells that fail mitosis, and anti-tumor actions of taxanes and MPS1 inhibitors, may depend on this effect.

## Introduction

Mitotic errors contribute to birth defects, aging, carcinogenesis and cancer therapy. They occur at a low frequency in normal cells, a higher frequency in cancer cells and a much higher frequency if mitosis occurs in the presence of chemotherapeutics (Fenech and Morley, 1985; Nelson et al., 2020). The genetic consequences of mitotic errors include structural rearrangements such as chromothripsis (Crasta et al., 2012) and numerical aberrations termed aneuploidy. A singular instance of structural or numerical defect can cause sustained genetic instability (Passerini et al., 2016; Umbreit et al., 2020). The cytological consequences of mitotic errors include micronuclei and chromatin bridges. Micronuclei exhibit abnormal nuclear transport and a high frequency of DNA damage in the subsequent cell cycle (Ly et al., 2017; Terradas et al., 2012). They are also prone to rupture which exposes their chromatin to the cytoplasm (Hatch et al., 2013). Chromatin bridges are a distinct type of mitotic error that are produced by dicentric chromosomes, merotelic attachments and catenations (Chan et al., 2007; Cimini et al., 2001; Maciejowski et al., 2015). They are typically resolved during anaphase (Chan et al., 2007) but can remain intact into the subsequent interphase when they become highly stretched due to tension from cell migration (Umbreit et al., 2020). Stretched chromatin bridges exhibit compromised nuclear envelopes, DNA damage and ultimately break through actin-mediated traction forces or endonuclease activity (Maciejowski et al., 2015; Umbreit et al., 2020). Broken chromatin bridges retract into the primary nucleus or become encapsulated into micronuclei (Hoffelder et al., 2004).

In addition to genetic and cytological consequences, mitotic errors induce inflammation and immunosurveillance through the activation of the viral DNA sensor, cGAS (Harding et al., 2017; Mackenzie et al., 2017). Upon binding to DNA, cGAS synthesizes 2’3′-cGAMP (cGAMP) which in turn activates STING followed by TBK1 and IRF3, ultimately leading to induction of type-1 interferon (IFN) expression and secretion (Chen et al., 2016). cGAS was originally proposed to discriminate viral from self-DNA by cytoplasmic localization of viral DNA (Sun et al., 2013). Additional regulatory mechanisms have now been identified, which include cGAS inhibition by nucleosomes and BANF1 as well as post-translation modifications (Guey et al., 2020; Kujirai et al., 2020; Pathare et al., 2020; Wu and Li, 2020). cGAMP can move between cells to activate STING in a paracrine manner in cell culture (Ablasser et al., 2013) and in tumors (Marcus et al., 2018). This may allow efficient signal propagation from cancer cells that have evolved blocks to interferon secretion. IFN activates adaptive immune responses, which makes cGAS activation an attractive therapeutic strategy to sensitize tumors to immune checkpoint inhibitors (Borden, 2019; Wang et al., 2017).

Anti-mitotic drugs (A-Ms) are a class of cancer chemotherapeutics that perturb mitosis and greatly increase the frequency of mitotic errors (Janssen and Medema, 2011). They are ideal tool compounds to investigate cGAS activation after mitotic failure because they induce chromosome missegregation in distinct manners that are independent of DNA damage. Taxanes are an important class of clinical A-Ms which stabilize microtubules and induce solid tumor regression. At saturating concentrations in cell culture, taxanes induce a prolonged mitotic arrest leading to cell death (Gascoigne and Taylor, 2008). Whether this mechanism is responsible for their tumor regression activity remains controversial (Komlodi-Pasztor et al., 2011; Weaver, 2014). Potent and specific A-Ms that target Aurora A Kinase, Aurora B Kinase, Polo-like Kinase 1 and KIF11 were tested in cancer patients but found to lack tumor-regression activity for reasons that remain unclear (Mitchison, 2012; Yan et al., 2020). We proposed that the special therapeutic activity of taxanes may depend on cGAS activation (Mitchison et al., 2017). Here, we test this idea by comparing the ability of different A-Ms to activate cGAS and correlating this with cytological defects. Unexpectedly, we found a key role for chromatin bridges in cGAS activation which could explain the higher clinical efficacy of taxanes.

## Results

### Co-culture Assay to Measure IFN Secretion

We developed a two-cell, co-culture assay to measure type-1 interferon (IFN) secretion caused by drug-induced mitotic failure in a multi-well plate format. The assay was designed to model paracrine communication between tumor cells and infiltrating leukocytes and report on physiologically relevant concentrations of secreted IFN. Because many cancer cell lines are defective in cGAS-STING-IFN signaling, we utilized an immortalized fibroblast line, hTERT-BJ 5Ta (BJ), where the pathway is known to be intact (Xia et al., 2016). To detect secreted IFN, we used THP1 monocytes which were engineered to express Lucia luciferase protein under the control of an IFN responsive promoter element (see methods). IFN producer cells, typically BJ fibroblasts, were treated with anti-mitotic drugs for 3 days to allow the accumulation of mitotic errors and activation of cGAS. We then seeded reporter THP1 monocytes for 18 hrs. to sense secreted IFN which we assayed with a plate-reader (Figure 1A). cGAS activation produces two paracrine signals, secreted IFN and 2’3′-cGAMP (cGAMP), which could activate the reporter cells. To focus the assay on secreted IFN, we utilized a THP1 reporter line in which STING was deleted using CRISPR technology. This line was non-responsive to direct cGAMP stimulation but did respond when cultured with BJ cells stimulated with cGAMP (Figure S1A). To confirm IFN secretion by BJ cells, we stimulated the cells with cGAMP and then stained for STAT1 phosphorylation, a canonical marker of IFN signaling (Darnell et al., 1994). We observed a dose-dependent increase in nuclear phospho-STAT1 signal following cGAMP treatment (Figure S1B), which confirms that cGAMP induces IFN secretion in the BJ cells and agrees with the co-culture assay result.

**Figure 1:**
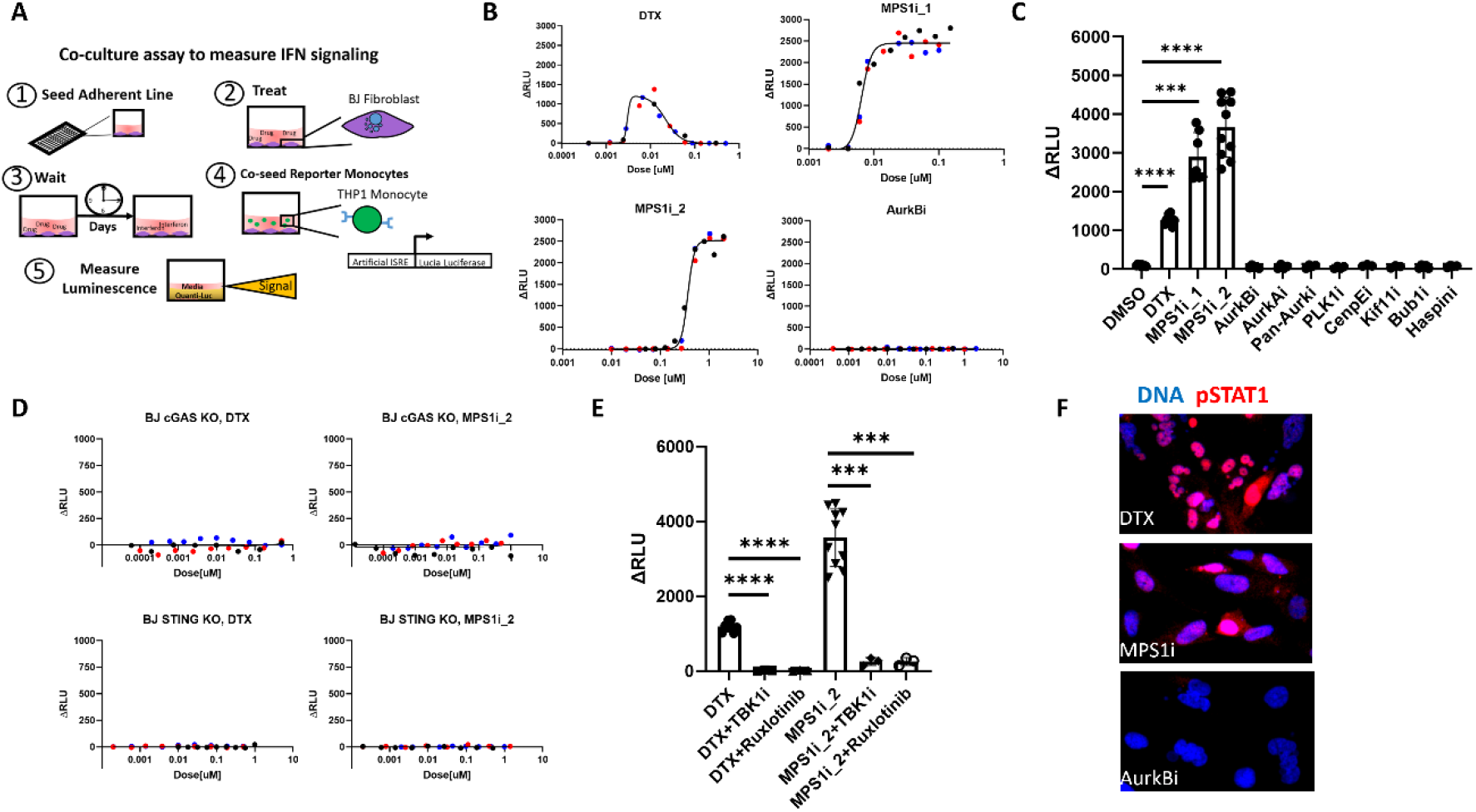
MT Stabilizers and MPS1 inhibitors induce IFN secretion through the cGAS-STING axis. **A:** Schematic of luciferase co-culture assay for measuring secreted IFN. The assay consists of 1) adherent line is seeded in a 96-well plate, 2) cells are exposed to drug, 3) cells are cultured in drug for multiple days, 4) reporter monocytes are co-seeded with the adherent lines for 18 hours and 5) the luciferase protein produced by the reporter monocytes, a proxy for the IFN secreted from the adherent line, is assayed with a plate reader. **B:** Dose-response for IFN signaling measured with the co-culture luciferase assay. Docetaxel (DTX) produces a bell-shaped curve centered around 10 nM. MPS1 inhibitors (MPS1i_1 and MPS1i_2) produce a hyperbolic response. AurkB inhibition (AurkBi) did not induce detectable IFN. Background luminescence measured from vehicle controls are subtracted. Different color markers represent independent experiments. See also Figure S2. **C:** IFN response measured across a panel of anti-mitotics. Doses used are found in Table 1. Only MT stabilizers and MPS1 inhibitors cause IFN signaling. Markers represent independent experiments. **D:** BJ cells lacking either cGAS or STING do not produce IFN in response to MT stabilizers and MPS1 inhibitors. Different color markers represent independent experiments. See also Figure S2. **E:** TBK1 inhibition and JAK 1/2 inhibition suppresses IFN induction by MT stabilizers and MPS1 inhibitors. Markers represent independent experiments. **F:** pSTAT1 increased expression and nuclear localization is observed after BJ cells are treated with DTX or MPS1i which cause IFN secretion but not after AurkBi.

We next tested whether the co-culture assay can detect drug-induced IFN secretion. We used the ATM inhibitor, KU-55933, that was previously reported to activate cGAS-STING signaling (Härtlova et al., 2015). We measured a dose-dependent increase in IFN signaling in the co-culture assay (Figure S1C). IFN induction by the ATM inhibitor was further confirmed by the nuclear localization of phospho-STAT1 in the fibroblasts (Figure S1D).

During assay development, we tested the response of multiple adherent cell lines from cancer and non-cancer backgrounds to exogenous cGAMP stimulation (Figure S1E-F). Only the three non-cancer lines tested responded with IFN production. BJ fibroblasts were the most sensitive and were therefore chosen for further assays. An important additional benefit of using non-transformed cells is that they reliably arrest in G1 following mitotic errors. We confirmed this arrest using EdU incorporation to measure progression to S-phase (Figure S1G-H). G1 arrest avoids complex errors due to multiple rounds of DNA replication and mitosis. cGAS activation in cancer-derived cell lines was investigated with a modified assay, as detailed below.

### MT Stabilizers and MPS1 Inhibitors Activate the cGAS-IFN Pathway

We assembled a panel of anti-mitotic drugs (Table 1) with disparate clinical efficacy (Penna et al., 2017; Rowinsky, MD, 1997; Traynor et al., 2011) in order to perturb mitosis in different ways and measure the corresponding IFN secretion. Only two drug classes generated robust and reproducible IFN responses after 2-3 days of exposure: the MT stabilizer, Docetaxel (DTX), and two MPS1 inhibitors, BAY1217389 (MPS1_1, 45) and CFI-402257 (MPS1i_2, 46) (Figure 1B). In dose-response assays, DTX produced a maximal IFN signal at ∼10 nM which is in line with estimates for intra-tumoral concentrations (Zasadil et al., 2014). DTX did not induce signal from reporter monocytes in the absence of fibroblasts (Figure S2A). Higher doses of DTX decreased IFN signal which we interpret to be due to cytotoxicity from prolonged mitotic arrest. We assayed viability with a standard readout of ATP and estimated the IC50 to be ∼50 nM which matches the reduction in IFN signal (Figure S2B). In time-response assays, the DTX-induced IFN signal peaked in intensity between 3 and 4 days in drug (Figure S2C).

**Table 1:** Overview of the Anti-mitotics Studied.

| Target | Drugs | Abbreviation<br>Used in Text | Clinical<br>Status | % Clinical<br>Response* | Primary<br>Dose<br>Studied<br>[nM] | IFN<br>Detected? |
| --- | --- | --- | --- | --- | --- | --- |
| MT<br>Dynamics | Docetaxel | DTX | First-line<br>therapy | Ovarian :<br>25-41%<br>NSLCS: 19-25%<br>Head and<br>Neck: 32-42% | 10 | Yes |
| MPS1 | BAY-1217389<br>CFI-402257 | MPS1i_1<br>MPS1i_2 | In Clinical<br>Trials | NA | 20<br>750 | Yes |
| Aurora B<br>Kinase | Barasertib | AurkBi | Failed<br>Trials | 0% | 250 | No |
| Aurora A<br>Kinase | Alisertib | AurkAi | Failed<br>Trials | 0%, 3%,<br>1% | 150 | No |
| Pan-Aurora<br>Kinase | Tozasertib | Pan-Aurki | Failed<br>Trials | 0% | 250 | No |
| Polo-like<br>Kinase 1 | Volasertib | PLK1i | Failed<br>Trials | 4.6%, 3.4%<br>,6.9% | 100 | No |
| Cetromere<br>Protein E | GSK-923295 | CenpEi | Failed<br>Trials | 3% | 250 | No |
| Kif11 | Ispinesib | Kif11i | Failed<br>Trials | 0%, 0% | 2.5 | No |
| Bub1 | BAY<br>1816032 | Bub1i | Pre-<br>clinical | NA | 1000 | No |
| Haspin | CHR-6494 | Haspini | Pre-<br>clinical | NA | 250 | No |

This is similar to the kinetics of accumulation of defective nuclear morphology observed in a mouse tumor model (Orth et al., 2011). Induction of IFN by MPS1 inhibitors followed a hyperbolic curve (Figure 1B). These drugs were significantly less toxic than the MT stabilizers (Figure S2D) and the IFN signal induced peaked after ∼2 days in drug (Figure S2E).

IFN induction by MT stabilizers and MPS1 inhibitors is likely to be an on-target activity by four criteria: 1) IFN was measured using multiple compounds with distinct chemical structures for each target (Figure S2F-G) 2) the shape of the dose-response curve was identical within each target class, though shifted on the concentration axis according to the potency of the drug, 3) DTX and MPS1 inhibitor EC_50_ values were similar for IFN induction and cell growth inhibition and 4) IFN induction by A-Ms decreased at high plating density, presumably due to reduced proliferation, while IFN induction by cGAMP was proportional to density (Figure S2I-J).

Surprisingly, all other A-Ms tested failed to produce measurable signal in our co-culture assay. In dose-response assays, the Aurora B kinase inhibitor (AurkBi), Barasertib, caused severe mitotic defects and nuclear abnormalities but failed to produce detectable IFN signal over 3 logs of concentration centered on the published IC_50_ (Figures 1B-C, 42).

We next verified that the IFN secreted after A-M exposure was induced by the cGAS-STING pathway. Fibroblasts where cGAS or STING were deleted using CRISPR engineering failed to produce IFN signal following treatment with MT stabilizers or MPS1 inhibitors (Figure 1D, Figure S2K). The luciferase signal induced by A-Ms was also suppressed by concurrent treatment with TBK1 or JAK1/2 inhibitors (Figure 1E). These results confirm that IFN induction by A-Ms is induced by cGAS-STING activation.

To confirm IFN induction with an alternative assay, we stained BJ cells for phospho-STAT1 after 3-day drug exposure and observed nuclear localization in response to DTX and MPS1i but not AurkBi (Figure 1F). This confirms the co-culture assay result.

Overall, our results suggest that IFN signaling is not a general consequence of mitotic failure but rather selectively induced through the cGAS-STING signaling axis by MT stabilizers and MPS1 inhibitors.

### Chromatin Bridges Correlate with cGAS Activation after Anti-mitotic Treatment

To understand why MT stabilizers and MPS1 inhibitors activate cGAS and other A-Ms do not, we quantified defects in nuclear morphology after A-M exposure. We established three classifications of nuclear abnormalities based on visual inspection: micronuclei, gross nuclei and chromatin bridges (Figure 2A, Figure S3A). Micronuclei were previously implicated in cGAS activation (Harding et al., 2017; Mackenzie et al., 2017). To score micronuclei we implemented an automated image analysis pipeline comprising of a pre-trained neural network coupled with a size threshold (Hollandi et al., 2020). A segmented image is shown in figure S3B. We then normalized the micronucleus count to the number of cells present within a given field. The resulting automated micronucleus scores agreed with manually scored images in a test set (Figure S3C). Gross nuclei and chromatin bridges were difficult to score algorithmically because they lacked stereotypical morphologies and were dimly stained with DNA dyes. To score these structures, we used visual inspection, scoring 3-5 images from multiple independent samples and normalizing to the number of cells in that field.

**Figure 2:**
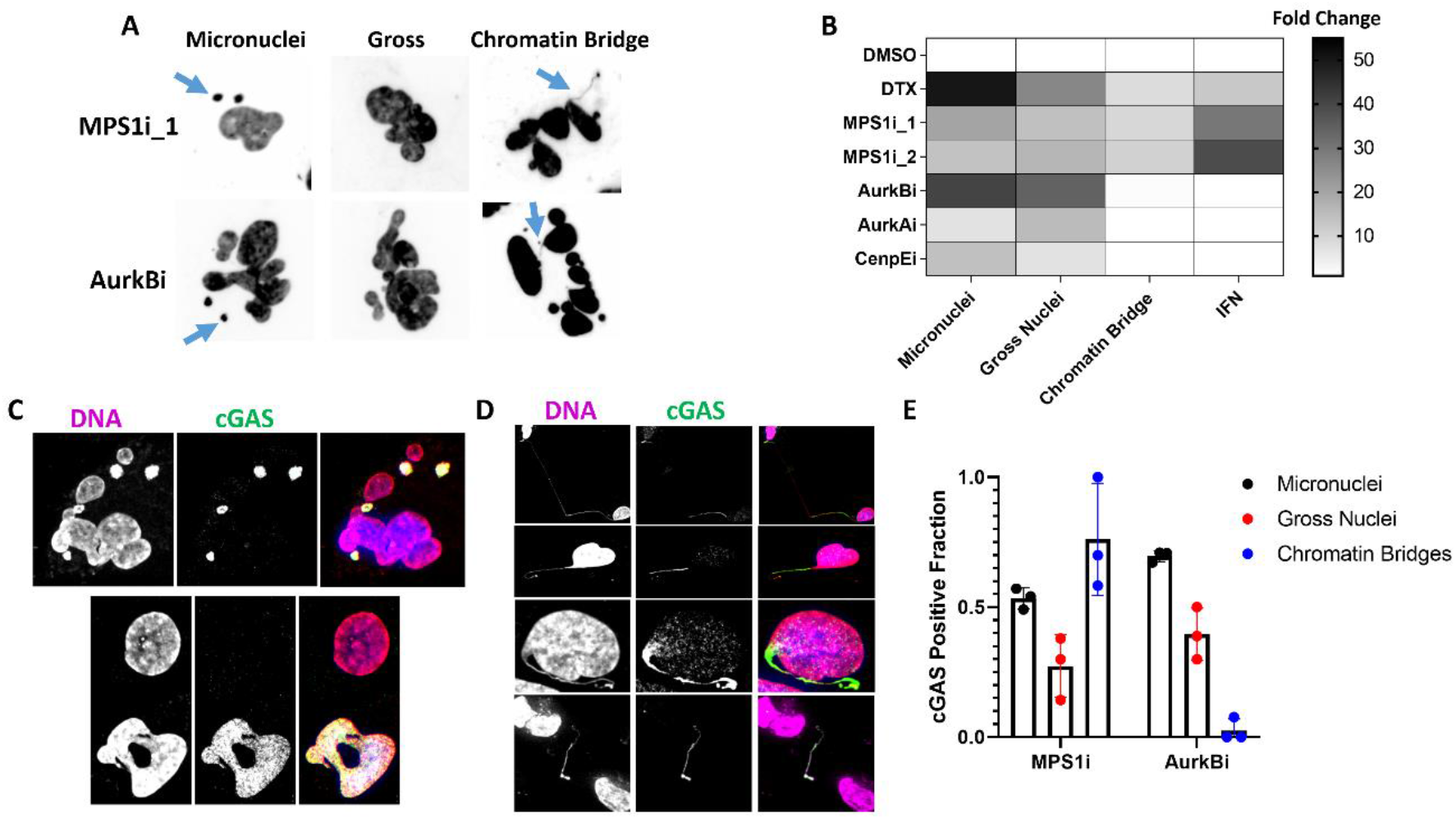
Quantification of aberrant chromatin structures and cGAS localization produced by anti-mitotics. **A:** Example nuclear morphologies produced by MPS1 inhibitors or Aurora B kinase inhibitors and visualized with a DNA dye. Aberrant structures are classified as micronuclei, gross nuclei or chromatin bridges. Example micronuclei and chromatin bridges are highlighted with blue arrows. See also Figure S2. **B:** A heatmap that summarizes the observed results for abnormal nuclear structures and IFN signal across different anti-mitotics. Chromatin bridges, not micronuclei, correlate with IFN signaling. Data was normalized to the corresponding vehicle control value. See also Figure S2. **C:** Images of cGAS positive micronuclei and gross nuclei produced after mitotic failure. cGAS was visualized with immunofluorescence. **D:** Examples of cGAS-positive chromatin bridges. Observed chromatin bridges were either intact (top row), broken on one end (second and third row) or broken on both ends (bottom row). cGAS was visualized with immunofluorescence. **E:** Quantified frequency of cGAS positivity on different types of aberrant chromatin structures produced after either MPS1 inhibition or AurkB inhibition. cGAS-positive micronuclei and gross nuclei showed similar frequencies after treatment with either drug. cGAS-positive chromatin bridges distinguish the two types of anti-mitotics. Each marker represents an independent experiment in which 3-5 FOVs were scored.

All tested A-Ms caused significant increases in micronuclei and gross nuclei (Figure S3D; 36). However, only DTX and MPS1 inhibitors caused significant increases of chromatin bridges (Figure S3D). Nuclear morphology scores and co-culture measurements of IFN signaling were combined into a heat map (Figure 2B). This presentation highlights our conclusion that chromatin bridges, not micronuclei or gross nuclei, correlated with IFN signaling after drug-induced mitotic failure.

### cGAS Localizes to All Aberrant Chromatin after Mitotic Errors

We next localized cGAS after mitotic errors by immunofluorescence of fixed cells (Figure S3E). To validate the cGAS antibody, we stained cGAS -/- BJ cells with mitotic errors and confirmed lack of signal (Figure S3F). In normal BJ cells, a significant fraction of interphase nuclei stained positive for cGAS. Micronuclei and gross nuclei induced by all drugs tested showed strong cGAS enrichment (Figure 2C). These observations show that cGAS localization to micronuclei is not a reliable proxy for cGAS activation, contrary to assumptions in recent papers (Gratia et al., 2019; Melms et al., 2020).

Chromatin bridges, which were only observed after treatment with MT stabilizers or MPS1 inhibitors, stained strongly for cGAS (Figure 2D). Interestingly, we observed distinct morphologies and staining patterns. Some chromatin bridges spanned two sister cells (2D; top panel) while others showed evidence of breakage (2D; middle two panels). Rarely we observed cGAS-positive chromatin bridges lacking any detectable connection to nuclei (Figure 2D; bottom panel).

We quantified the frequency of cGAS localization to all abnormal chromatin structures after either MPS1 inhibition or AurkB inhibition (Figure 2E). cGAS localized to micronuclei and gross nuclei to a similar degree in both, while cGAS-positive chromatin bridges were only observed after MPS1 inhibition.

We tried to quantify the cGAS:DNA ratio for different aberrant structures. Chromatin bridges stained strongly for cGAS, while the density of DNA in bridges, based on DNA dye fluorescence signal per unit area, was typically very low. This low DNA signal precluded accurate measurement of the cGAS:DNA ratio in bridges but we are confident that it is higher, and probably much higher, than in micronuclei, gross nuclei or control nuclei. Our results suggest that all chromatin can recruit cGAS. Aberrant nuclear structures (micronuclei and gross nuclei) formed after mitotic errors recruit more cGAS than normal nuclei but do not activate it. Chromatin bridges recruit the highest amount of cGAS per unit DNA and only they activate it sufficiently to induce IFN after failed mitosis.

### cGAS Activation Requires Cytokinesis and Stretching Forces on Chromatin

We next tested whether blocking chromatin bridge formation would supress IFN signal after failed mitosis. We chose to induce chromatin bridges with MPS1 inhibitors because they are minimally cytotoxic in a 3 day assay (Figure S2D). MPS1 inhibitors also reduce the short-term cytotoxicity of other A-Ms because they compromise the spindle asssembly checkpoint which prevents cell death from prolonged mitotic arrest (Figure S2H, 47). We treated cells with a constant concentration of MPS1i sufficient to activate cGAS combined with a titration of different A-Ms (Figure 3A). These drugs were chosen to block cytokinesis by different mechnaisms, with the exception of CenpE inhibition which is not known to block cytokinesis (Tanudji et al., 2004) and was considered a negative control. In brief, inhibition of AurkA and KIF11 cause monopolar spindles, inhibition of AurkB, Bub1 and Haspin prevent spindle midzone assembly and inhibiton of MT polymerization prevents spindle and midzone formation (Tischer and Gergely, 2019). We also inhibited F-actin which blocks cytokinesis by preventing furrow contraction as well as actin-dependent traction forces that move daughter cells apart after mitosis (Figure 3B, Copeland, 1974). All drugs that interfered with cytokinesis blocked IFN induction by MPS1i. In contrast, the CenpE inhibitor did not block IFN signal. IC_50_ values for the reduction of IFN signal were consistent with on-target activity for all compounds. The results for AurkB and AurkA inhibition was confirmed with structurally dissimilar drugs.

**Figure 3:**
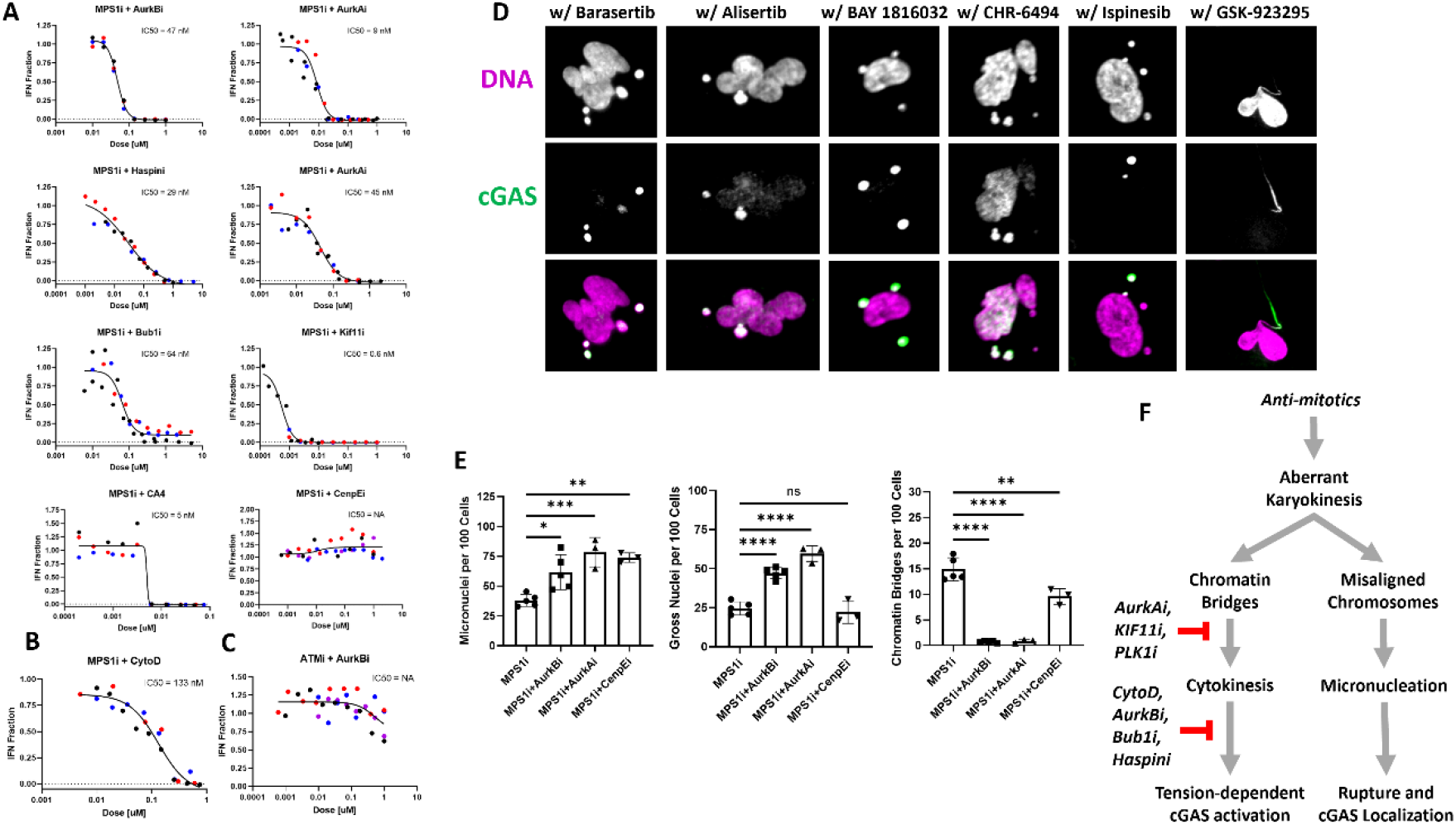
cGAS activation requires cytokinesis and stretching forces on chromatin. **A:** The effect of anti-mitotic combinations on IFN signal. The majority of tested anti-mitotics show dose-dependent suppression of the IFN signal produced by MPS1 inhibition. CenpE inhibition is not known to interfere with cytokinesis and is considered a negative control. Values are reported as the percent IFN signal relative to the MPS1 inhibition alone. Different color markers represent independent experiments. **B:** Low-dose actin inhibition by Cytochalasin-D suppresses IFN produced by MPS1 inhibition. Different color markers represent independent experiments. **C:** AurkB inhibition does not suppress IFN signal generally. DNA damage caused by ATM inhibition activated cGAS-STING-IFN axis irrespective of AurkBi dose. Different color markers represent independent experiments. **D:** Example images of cGAS-positive micronuclei observed after anti-mitotic combinations. Notably, only CenpE inhibition permits cGAS-positive chromatin bridge formation when combined with MPS1 inhibition. This combination also retains IFN signaling. **E:** Frequencies of aberrant chromatin structures produced by anti-mitotic combinations. Anti-mitotic combinations which suppress IFN signal in panel A also suppress chromatin bridge formation. Markers represent independent experiments. **F:** Model for how anti-mitotics affect mitotic failure and IFN signal. Only MT stabilizers and MPS1 inhibitors produce chromatin bridges because they do not overtly interfere with spindle formation or cytokinesis. All anti-mitotics tested produce cGAS-positive micronuclei. Chromatin bridge formation is directly perturbed by many anti-mitotics through targeting spindle formation (AurkAi and KIF11) or cytokinesis (Cytocholasin D and AurkBi) or both (CA4).

It is possible, although unlikely, that these drugs supressed cGAS-STING signaling through an unknown mechanism or by off-target inhibition of a signaling pathway component such as TBK1. We tested the AurkBi by combining it with an ATM inhibitor, KU-55933, which activates cGAS in a mitosis-independent manner (Figure S1C, 44). The AurkBi had no effect on IFN signaling produced by ATM inhibition (compare Figures 3C to top left of 3A). We conclude that the ability of the AurkBi to supress IFN induction by the MPS1i is due to on-target interference with cytokinesis.

We next performed immunofluorescence to visualize cGAS localization in cells treated with drug combinations. Most combinations induced high levels of cGAS-positive micronuclei irrespective of whether they induced IFN secretion (Figure 3D). In contrast, cGAS-postive chromatin bridges were only seen in the conditions where IFN was induced, MPS1i + CenpEi (Figure 3D; far right). We quantified the frequency of different chromatin structures in a subset of drug combinations and observed that supression of chromatin bridges corresponded to inhibition of IFN signaling (Figure 3E). These data show that cytokinesis is required for cGAS activation after mitotic errors and also provide additional evidence that cGAS-positive micronuclei do not drive cGAS activation and IFN signaling. In principle, chromatin bridges can be formed in the absence of cytokinesis but will remain unstretched (Fenech, 2006). We made efforts to analyze chromatin bridges in cells where cytokinesis was blocked, but dense packing of chromatin in these cells made scoring of thin strands difficult. Thus, our data do not formally distinguish chromatin bridge formation from chromatin bridge stretching (discussed below).

Based on our data, we propose the model of how A-Ms activate cGAS diagrammed in Figure 3F. Aberrant chromsome segregation is enhanced by all anti-mitotics but only DTX and MPS1 inhibition cause chromatin bridges that activate cGAS.

### Paracrine cGAMP assay to measure cGAS activation in Cancer Cells

To generalize our results, and test their relevance for drug action in tumors, we sought to measure cGAS activation by mitotic errors in cancer cells. Many cancer lines are unable to secrete IFN in response to aberrant DNA or cGAMP (32, 51, Figures S1E-F). Consistent with these reports, two cancer lines, MDAMB231 and HeLa, failed to produce detectable IFN secretion after treatment with either DTX or MPS1i (Figure 4A). We therefore developed a cGAS activation assay that measures paracrine spread of cGAMP from cancer cells to cGAS -/- BJ fibroblasts, which then secrete IFN that is detected by reporter STING -/- THP1 monocytes (Figure 4B). As we tested different cancer cell lines in this assay, we detected large differences in basal cGAMP production (Figure S4A). MDAMB231 cells, which are known to have chronically activated cGAS (Bakhoum et al., 2018), exhibited the largest basal signal in our tri-culture assay. We confirmed that co-culture of MDAMB231 cells with BJ fibroblasts activated IFN secretion by staining for phospho-STAT1 (Figure S4B). Importantly, co-cultures of cancer lines with STING -/- BJ cells, or in the presence of a TBK1 inhibitor, failed to produce detectable IFN which validates the assay principle (Figure S4C). The three-cell assay provides a sensitive and convenient technique to detect cGAS activity in cancer lines which lack IFN secretion and may model the tumor microenvironment.

**Figure 4:**
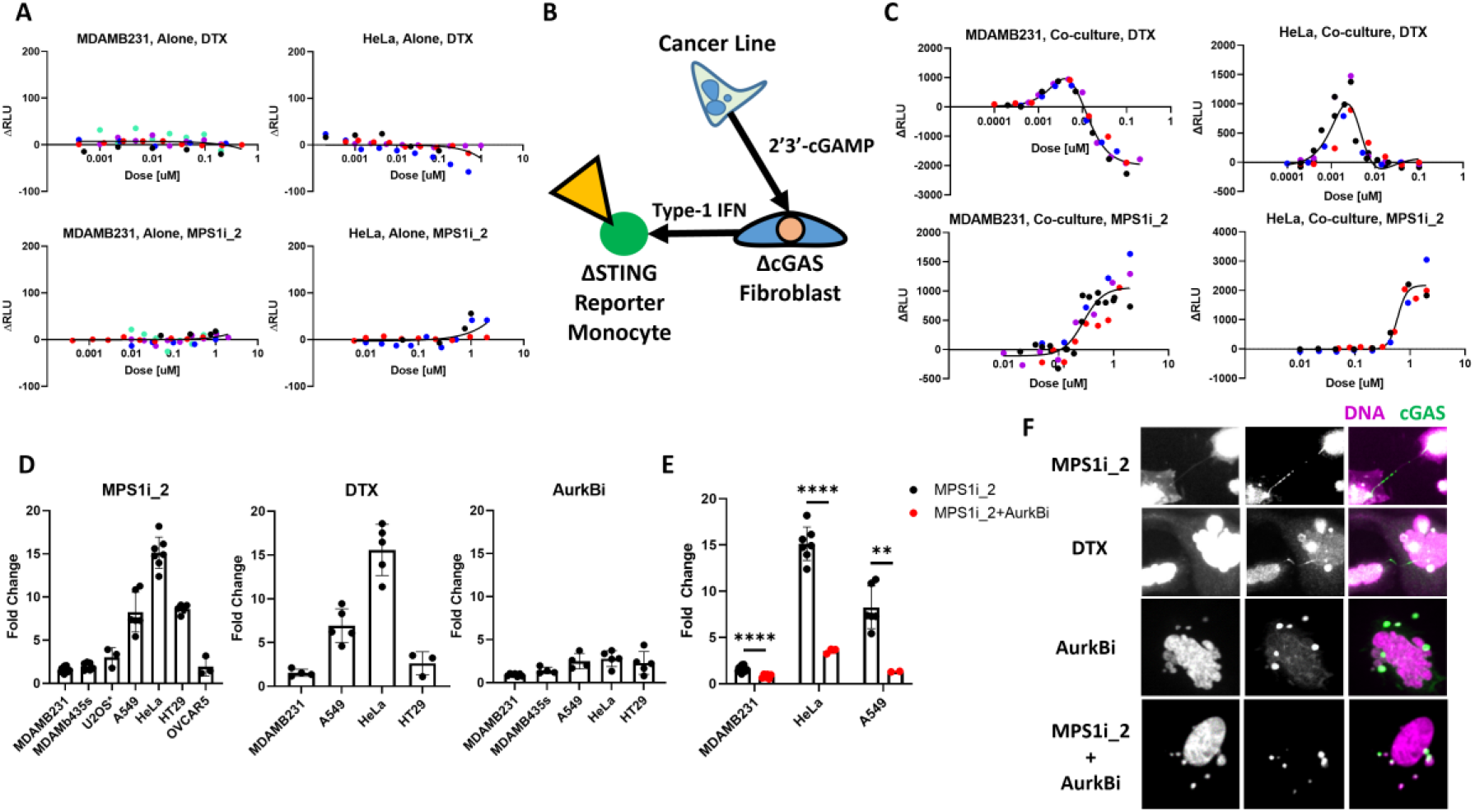
Anti-mitotics activate cGAS in cancer lines through chromatin bridge formation. **A:** Cancer lines do not produce IFN in response to MPS1i or DTX. Different color markers represent independent experiments. **B:** Schematic of the modified co-culture luciferase assay designed to detect cGAS activation in cancer cells through paracrine cGAMP signaling **C:** MPS1 inhibitors and MT stabilizers activate cGAS in cancer lines. Both MDAMB231 and HeLa cells showed dose-dependent responses to DTX and MPS1 inhibition. These response curves mirrored the shape of IFN induction in the BJ cells. The negative ΔRLU in the DTX MDAMB231 response represents cytotoxicity. Different color markers represent independent experiments. **D:** MPS1 inhibitors and MT stabilizers activate cGAS across a panel of cancer cells. 1 nM DTX, 750 nM MPS1i_2 or 250 nM AurkBi were used. Cancer lines showed variable levels of response to anti-mitotic treatment. Notably, of the cancer lines that did show cGAS activation, MPS1 inhibition and DTX treatment induced larger responses than AurkBi. Markers represent independent experiments. **E:** AurkB inhibitors reduce cGAS activation in MPS1i treated cells. Markers represent independent experiments. **F:** HeLa cells treated with different anti-mitotics. cGAS-positive chromatin bridges are produced after treatment with MPS1i and DTX, but not by AurkBi or the combo AurkBi+MPS1i. Notably, conditions which exhibit minimal cGAS activation contain abundant cGAS-positive micronuclei.

### Chromatin bridges, not micronuclei, activate cGAS in cancer cells

We next used the three-cell assay to measure cGAS activation in response to anti-mitotic treatment. We selected two cancer lines to test the role of chromatin bridges, MDAMB231 which exhibited high basal cGAS activity and HeLa which exhibited low basal activity (Figure S4A). DTX and MPS1 caused cGAS activation with dose-response curves similar to those measured in BJ cells (Figure 4C, compare to Figure 2B). Negative ΔRLU values represent cytotoxicity from drug that reduced luminescence below the basal signal. We then treated a collection of different cancer cells with A-Ms at on-target doses (Figure 4D). Cancer lines displayed variable levels of signal induction, reported as fold change relative to their basal signal, in response to DTX and MPS1 inhibition. Notably, lines which were highly responsive to MPS1i and DTX only showed minimal signal after AurkB inhibition. Inhibition of AurkB blocked the additional cGAS activation induced by MPS1i in multiple lines, confirming a requirement for cytokinesis in cGAS activation after mitotic failure (Figure 4E). We then performed immunofluroescence to determine cGAS localization. Most cancer lines exhibited constituitive levels of abnormal nuclear structures that stained positive for cGAS (Figure S4D). We stained HeLa cells after treatment with different anti-mitotics (Figure 4F). Similar to our results with fibroblasts, cGAS was found to localize to micronuclei, gross nuclei and chromatin bridges, but cGAS activation was exclusively associated with the presence of cGAS-positive chromatin bridges with very high cGAS:DNA staining ratios.

Similar results were observed in MDAMB231 cells (Figure S4E). Thus, the model presented in Figure 3F applies to cancer lines, with the complication that cGAS activation by chromatin bridges is superimposed on varying levels of constituitive cGAS activation.

## Discussion

We studied a panel of A-M drugs that produce mitotic errors through distinct mechanisms and measured the corresponding amount of IFN secretion in immortalized fibroblasts and cancer cell lines. Unexpectedly, we observed that chromatin bridges, not micronuclei, activate cGAS after a failed mitosis in human cells despite both structures recruiting cGAS.

Our conclusion that abundant cGAS-positive micronuclei generated by A-Ms do not activate cGAS in human cell lines appears to contradict previous reports (Harding et al., 2017; Mackenzie et al., 2017). Careful microcopy is required to reliably image and score thin chromatin bridges and they might have been missed or underestimated in previous studies. It is also possible that micronuclei formed in other ways, or in other species, can activate cGAS, e.g. following irradiation of mouse cells. cGAS activation after irradiation is difficult to ascribe solely to mitotic errors because it simultaneously inflicts interphase DNA damage, oxidative stress and gene upregulation which combine to modulate cGAS activation in complex ways (Vanpouille-Box et al., 2017). IFN signaling detected 5-7 days post-irradiation could be an indirect effect of chromatin bridges breaking and becoming micronuclei or micronuclei incurring massive DNA damage in subsequent cell cycles (Ly et al., 2017). Our work used mitosis-specific perturbations and a cell line in which mitotic errors cause G1 arrest, which prevents the complex outcomes seen in cells undergoing multiple cell cycles with damaged chromatin. Species differences in cGAS activity may also contribute to differences in cGAS activation by micronuclei. DNA damage in micronuclei is likely to generate short DNA fragments. Human cGAS is minimally activated by short fragments, while mouse cGAS is strongly activated (Zhou et al., 2018). Further studies are required to clarify cGAS activation by micronuclei in different species.

To reliably quantify cGAS activation, we developed two novel assays that are affordable and amenable to high-throughput screening. They were designed to mimic the tumor microenvironment by reporting on paracrine signaling between cancer, stroma and leukocytes. Our two-cell assay (Figures 1-3) used STING -/- reporter monocytes to measure physiologically relevant levels of IFN secreted by a non-transformed fibroblast line that may model tumor stroma. Our three-cell assay (Figure 4) used cancer cells, cGAS-/-fibroblasts and STING -/- reporter monocytes to measure paracrine spread of cGAMP from cancer cells to fibroblasts. This assay confirmed literature reports of strong basal activation of cGAS in some cancer lines (Bakhoum et al., 2018; Carozza et al., 2019) and efficient cell-to-cell spread of cGAMP (Ablasser et al., 2013). We did not investigate the mechanism of cell-to-cell spread, which was reported to occur via gap junctions (Ablasser et al., 2013) and also by transporter-mediated secretion-reuptake (Ritchie et al., 2019). Paracrine spread of cGAMP from cancer cells to stroma and leukocytes may allow the whole tumor to mount an IFN response to drug-induced mitotic errors even if the cancer cells have lost their ability to secrete IFN, which is often observed in cancer cell lines.

Our data suggests that among the A-Ms tested, MPS1 inhibitors and low, clinically-relevant concentrations of MT stabilizers produce chromatin bridges which activate cGAS. An important caveat is that we could only visualize chromatin bridges reliably when anaphase and cytokinesis were sufficiently successful that most chromatin was divided between two distinct daughter cells. Under conditions where spindles were monopolar or cytokinesis failed completely, aberrant chromatin accumulated in a large mass that we could not resolve by microscopy. The chromatin bridges we scored were almost always stretched or broken by forces from furrow ingression and post-mitotic actin-mediated traction motility (Umbreit et al., 2020). We do not know if some drugs, such as AurkB inhibitors, failed to induce chromatin bridges or blocked their stretching.

Our study leaves two important mechanistic questions unanswered: how are cGAS-activating chromatin bridges induced by MT stabilizers and MPS1 inhibitors and how do they activate cGAS? Based on literature review, we speculate that chromatin bridges are the product of unresolved catenations. Sister chromatids enter mitosis heavily catenated in both cancer and normal cells (Chan et al., 2007; Oliveira and Nasmyth, 2010). MPS1 inhibitors may interfere with decatenation by shortening the duration of metaphase, while MT stabilizers at low concentrations reduce tension across sister chromatids that promotes resolution. A small number of catenations persists into anaphase under normal conditions and are resolved by a PICH-BLM-TopoII system (Biebricher et al., 2013; Chan et al., 2007). We suspect that MT stabilizers and MPS1 inhibitors generate sufficient anaphase catenations to overwhelm the resolution machinery.

Stretched chromatin bridges recruited cGAS to a higher local density than other aberrant forms of post-mitotic chromatin. We hypothesize that tension plays two roles in cGAS recruitment and activation: displacement of nucleosomes and alignment of pairs of DNA duplexes side-by-side which facilitates cGAS oligomerization. Nucleosomes allosterically inhibit cGAS (Boyer et al., 2020; Kujirai et al., 2020; Pathare et al., 2020). They can be displaced by stretching forces in the low 10s of pN (Bustamante et al., 2003), which is well within the range that can be generated by furrow ingression and traction forces (Umbreit et al., 2020). The recently proposed “ladder model” of cGAS binding and activation may be directly relevant to cGAS activation by chromatin bridges (Andreeva et al., 2017). The optimal DNA configuration for cGAS activation is two parallel duplexes of naked DNA which promotes the assembly of 2:2 cGAS:dsDNA complexes wherein cGAS dimers are the rungs of the ladder. Stretching of catenated sister chromatids is expected to generate exactly this configuration in the form of two stretched hairpins on each side of the catenation.

It is not safe to extrapolate directly from cell culture data to clinical activity, but if we provisionally accept the hypothesis that IFN secretion contributes to tumor regression, then our data have interesting implications. They suggest taxanes promote tumor regression in part by inducing IFN after failed mitosis, and the clinical failure of high-quality inhibitors targeting Kif11, AurkA, AukB and Plk1 may be due to lack of IFN induction. A major caveat to this hypothesis is that most cancer cell lines are unable to secrete IFN in response to MT stabilizers or MPS1 inhibitors. It is possible that taxane-responsive solid tumors retain IFN secretion or that cGAMP spreads from tumor cells to neighboring stroma cells and leukocytes to activate STING. An alternative hypothesis, not mutually exclusive, is that taxanes promote an IFN response in proliferating stromal cells. These ideas could be tested by measuring IFN induction and other inflammatory cytokines in taxanes-treated tumors at single-cell resolution. An exciting implication of our data is that MPS1 inhibitors hold clinical promise because they efficiently activate cGAS. However, they are significantly less cytotoxic than taxanes in our short-term assays, which makes clinical efficacy hard to predict.

Lastly, our results have implications for immunosurveillance of failed mitosis in normal tissues. Mitotic errors occur spontaneously and can promote tumor development. cGAS activation by stretched chromatin bridges may be an evolved mechanism to alert the immune system to potentially transformed cells. Conversely, the same mechanism could promote tumorigenesis by generating inflammatory niches. Further work is needed to understand immunosurveillance of failed mitosis *in vivo*, and the roles of cGAS and cGAMP in tumor evolution.

## Acknowledgments

The authors would like to thank Yukiye Koide and Emma Spady from the Silver lab for assistance with CRISPR KO cell line generation, Mirra Chung from the Sorger Lab for sharing the MDAMB231 cell lines, the Nikon Imaging Center at Harvard Medical School, the Laboratory of Systems Pharmacology and Dr. Jon Kagan for reviewing the manuscript. P.F. was supported by a fellowship from the Lynch foundation. This work was funded by NIH grant GM131753.

## Author Contributions

Conceptualization, P.F., P.D., T.M.; Methodology, P.F., P.D., T.M.; Investigation, P.F.; Writing – Original Draft, P.F.; Writing – Review & Editing, P.F., P.D., T.M.; Funding Acquisition, P.F., T.M.; Resources, T.M.; Supervision, T.M.

## Declaration of Interests

The authors declare no competing interests.

**Table 1 Anti-mitotics studied:** The table describes the drug targets, abbreviations, clinical status and reported clinical response rates. Percent clinical response refers to objective response as a monotherapy in solid tumors (Penna et al., 2017; Rowinsky, MD, 1997; Traynor et al., 2011). The primary doses used refer to doses used in our experiments when dose-responses are not shown. These doses are based on literature and our own observations. IFN detected refers to whether we observed IFN signal in the co-culture assay.

## Materials and Methods

### Cell culture

Solid tumor lines were purchased from ATCC with the exception of MDAMB231 which was kindly provided by the Sorger lab at Harvard Medical School. The U2OS line used expressed fluorescently labeled cGAS and BAF and was kindly shared by Dr. Tae Yeon Yoo. hTert-BJ-5TA and hTert-BJ-5TA cGAS KO were kindly provided by the Silver lab at Harvard Medical School. Engineered THP1 reporter cells were purchased from Invivogen. All the cell lines used in this study were grown under 37°C and 5% CO_2_ in a humidified incubator. Cell lines were grown in appropriate media supplemented with 10% fetal bovine serum (FBS) and 1% (*v:v)* penicillin/streptomycin (pen/strep). Specifically, THP1, MDAMB231, OVCAR5 and HT29 cells were grown in RPMI; BJ-5Ta, U2OS, HeLa, MDAMB435s and A549 cells were grown in DMEM. Cells were kept below passage 25 and regularly screened for mycoplasma.

### Drugs

MT Stabilizers (Docetaxel; MedChemExpress, Epothilone-B;MedChemExpress), MT Depolymerizers (Combetastatin-A4, provided by Dr. Sergine Brutus), Aurora Kinase A Inhibitors (Alisertib;MedChemExpress, MK-8745;MedChemExpress), Aurora Kinase B Inhibitors (Barasertib; MedChemExpress, ZM-447439;MedChemExpress), Pan-Aurora Kinase Inhibitors (Tozasertib; Haoyuan chemexpress), MPS1 Inhibitors (CFI-402257;MedChemExpress, BAY-1217389;MedChemExpress), Haspin Inhibitor (CHR-6494; Cayman Chemicals), Bub1 Inhibitor (BAY-18116032;MedChemExpress), Eg5 Inhibitor (Ispinesib; Cayman chemicals), Centromere Protein E Inhibitor (GSK923295; Cayman Chemicals), Actin Depolymerizer (Cytocholasin-D; Provided by Dr. Christine Field), ATM Inhibitor (KU-55933;SelleckChem), TBK1 Inhibitor (MRT67307;Cayman Chemicals), JAK Inhibitor (Ruxlotinib;MedChemExpress). All drugs were dissolved in DMSO.

### Antibodies and DNA Stains

Anti-cGAS 1:400 (D1D3G;CST), Anti-phospho-STAT1 1:400 (Tyr701, 58D6;CST); secondary antibodies were purchased from Invitrogen and conjugated with different AlexaFluors (1:500). Hoechst 33528 (1-10 ug/mL) or Syto DeepRed (1:1000, ThermoFisher) were used for counterstaining DNA.

### Co-culture Reporter Assay

Assay is based on reporter cell protocol outlined by Invivogen, but modified to incorporate two cell lines, one IFN producer (i.e. BJ Fibroblasts) and one IFN reporter (i.e. THP1). Producer lines were resuspended in fresh media and seeded with 100 uL volume in a 96 well plate between 8-10,000 cells per well. Cells were treated with compound 6 to 24 hours after initial seeding using a D300 digital dispenser and all wells were normalized to same total volume of solvent, typically DMSO, below 0.1% total volume. Reporter cells were resuspended in fresh media and added to the drugged cells 72 hours after drug addition. Reporter cells were seeded with 100,000 cells per well in a volume of 50 uL and left overnight. To assay for luciferase secretion, 10 uL of conditioned media from the co-culture wells were mixed with 50 uL of the commercial Quanti-Luc reagent (Invivogen) in a black, transparent 96-well plate (Corning) and brought to a plate reader (Victor Plate Reader) to measure luminescence. Experimental conditions were typically performed with technical replicates in triplicate.

### Tri-culture Reporter Assay

Assay performed in an identical manner to the co-culture assay, but producer cells consisted of BJ cGAS KO fibroblasts mixed with cancer lines at a 1:1 ratio, 5000 cells each. The rest of the assay proceeded as described for the co-culture assay.

### Immunofluorescence

Cells were grown on #1.5 glass coverslips in multi-well plates (Corning) and exposed to drug. Following multi-day drug exposure, cells were washed with 37C degree PBS and fixed with either 37C degree 3.2% PFA (diluted in PBS) for 10 minutes at room temperature for cGAS staining or with ice-cold methanol at −20 for 10 minutes when for pSTAT1 staining. Coverslips were then washed with PBS 3x times and blocked and permed with PBS + 0.1% Tritonx100 +5% BSA for 1 hour at RT. Coverslips were then stained with primary antibodies suspended in PBS+0.1% Tritonx100 + 1% BSA overnight at 4C degrees. Coverslips were then brought to RT and washed 3x times with PBS+0.1% Tritonx100 and stained with secondary anti-bodies for 1 hour at RT. Coverslips were then washed 3x times with PBS+0.1% Tritonx100 and stained with DNA dye for 30 minutes. Coverslips were washed 3x times and mounted in media comprised of 20 mM Tris at pH 8, 0.5% N-propyl gallate and 90% Glycerol. Coverslips were sealed with nail-polish hardener (Sally Henson).

### EdU incorporation and detection

EdU (5-ethynyl-2′-deoxyuridine) detection was performed with the Click-iT EdU Alexa Fluor 555 Imaging Kit (Thermo-Fisher Scientific). EdU was diluted in DMSO to a final concentration of 10 mM. EdU was added to cell media following drug treatment to a final concentration of 10 uM. Cells were fixed 3.2% PFA for 10 minutes at room temperature. Detection of EdU-DNA was performed according to the Click-iT EdU Alexa Fluor 555 Imaging Kit per the manufacturer’s instructions.

### Imaging

Slides were brought to the microscope for imaging. 20x images were collected on a Nikon Eclipse Ti2-E inverted microscope equipped with Nikon CFI Plan Apo Lambda 20x, NA 0.75 objective lens. Images were acquired with a SOLA SE V-nIR light engine, and with an Andor Zyla 4.2 PLUS sCMOS camera and controlled with NIS Element software. 60x images were collected on a Confocal Nikon Ti inverted microscope equipped with Nikon CFI Plan Apo Lambda 60x, NA 1.4 objective lens. Z-series were acquired with a Spectral Applied Research LMM-5 laser module with solid state lasers and collected with a Hamamatsu ORCA-R2 cooled CCD camera. MetaMorph software was used for image acquisition.

### Automated scoring of nuclei and micronuclei

We developed an analysis pipeline for the automated segmentation and analysis of nuclei and micronuclei in images. Images were pre-processed with the ImageJ algorithms unsharp mask and fast Fourier transform band pass filter to improve the detection of micronuclei. The images were then uploaded to the publicly available nucleAIzer segmentation tool (Hollandi et al., 2020). Post-processing of the nucleAIzer output was performed in MATLAB to distinguish nuclei from micronuclei and to reduce over-segmentation of multi-lobed nuclei. Example results are shown in supplementary figure 3. Micronuclei frequencies were normalized by the number of cells scored in the same FOV and reported as # of micronuclei per 100 cells.

### Manual scoring of gross nuclei and chromatin bridges

Images were manually scored for gross nuclei and chromatin bridges. Gross nuclei were classified based on multiple factors including binucleation, multinucleation, deranged morphology and abnormally large size. Chromatin bridges were identified based on obvious DNA spanning two nuclear objects. However, due to the fragile nature of chromatin bridges, bridge fragments and the remnants of bridge protrusion were also included in our classification. Between 3-5 FOVs were analyzed per independent experiment and the frequency of abnormality was normalized to the total number of cells.

### CRISPR KO Cell Line Generation

Knockout cell lines were generated with the Synthego KO Kit (Redwood City, California). Single clones were expanded, sequenced and KO efficiency was estimated with the online Synthego ICE tool. Western blots were performed on selected clones to confirm STING KO (Figure S2).

### Statistical tests

Unpaired t-tests were used to compute reported P-values. When more than two conditions are compared, a one-way analysis of variance was used. Prism software (Graphpad) was used for all calculations. The level of statistical significance is represented as follows: n.s.=P>0.05, *=P≤0.05, **= P≤0.01, ***=P≤0.001 and ****=P≤0.0001.

