## Supplemental for "Chromatin Bridges, not Micronuclei, Activate cGAS after Drug-induced Mitotic Errors in Human Cells"

### Supplemental Data:

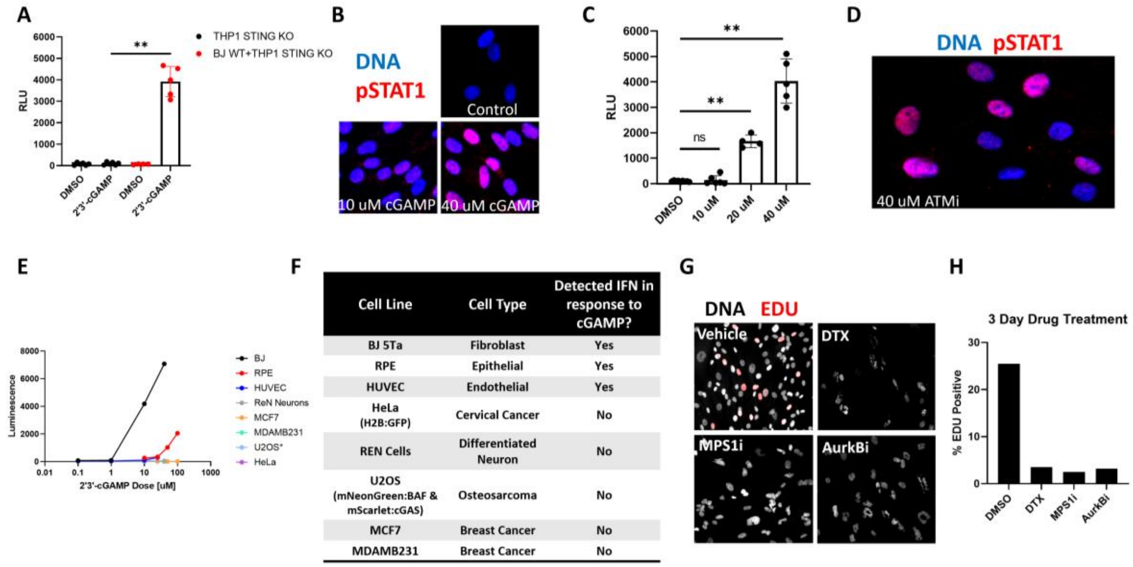


| Cell Line | Cell Type | Detected IFN in response to cGAMP? |
|---|---|---|
| BJ 5Ta | Fibroblast | Yes |
| RPE | Epithelial | Yes |
| HUVEC | Endothelial | Yes |
| HeLa (H2B:GFP) | Cervical Cancer | No |
| REN Cells | Differentiated Neuron | No |
| U2OS (mNeonGreen:BAF & mScarlet:cGAS) | Osteosarcoma | No |
| MCF7 | Breast Cancer | No |
| MDAMB231 | Breast Cancer | No |

**Supplemental Figure 1: Co-culture Assay Development**

**A:** THP1 STING KO reporter cells cultured with or without hTert-BJ-5Ta fibroblasts (BJ) were stimulated with 2'3'-cGAMP (cGAMP) for 18 hours and then assayed. THP1 STING KO cells did not produce luciferase in response to cGAMP when cultured alone but did produce detectable luciferase when co-seeded with cGAMP-stimulated BJ cells. Markers represent independent experiments.

**B:** BJ cells stimulated with cGAMP for 18 hours and stained for phospho-STAT1 (Y701). Increased expression and nuclear localization in response to cGAMP suggests intact STING-IFN signaling in the BJ cells and agrees with co-culture result.

**C:** Co-culture assay performed with the ATM inhibitor, KU-55933. BJ cells were treated with drug for 3 days prior to THP1 STING KO reporter co-seed. Markers represent independent experiments.

**D:** BJ cells show pSTAT1 nuclear translocation after 3 day ATM inhibition, which suggests activated cGAS-STING-IFN signaling and agrees with the co-culture assay results.

**E:** Dose-response for cell-lines tested in response to 2'3'-cGAMP stimulation. Non-transformed cell lines BJ, RPE and HUVEC showed variable responsiveness. BJ fibroblasts showed marked sensitivity to cGAMP. All cancer lines tested did not produce detectable IFN. Markers represent the average of technical replicates.

**F:** Table of cell lines tested in the co-culture assay with direct 2'3'-cGAMP stimulation.

**G:** BJ cells treated with anti-mitotics for 3 days and then pulsed with 10 uM EdU for 1.5 hours. Incorporated EdU was visualized.

**H:** Quantification of EdU incorporation after anti-mitotic treatment. Value bars represent the average frequency across 3-5 FOVs.

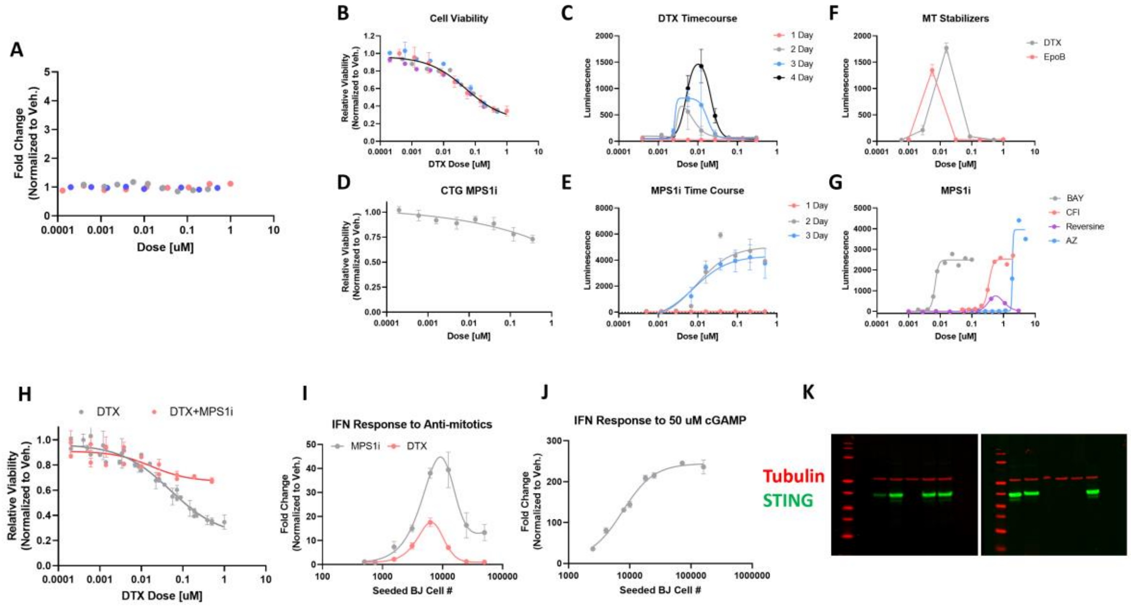


**Supplemental Figure 2: Anti-mitotics Activate cGAS-STING Signaling**

**A:** THP1 reporter cells exposed to DTX overnight do not exhibit reporter activation. Different colored markers represent independent experiments.

**B:** Cytotoxicity of DTX after 3 day treatment of BJ fibroblasts measured by ATP amount with Cell-titer Glo assay. The fitted IC50 was ~ 50 nM which matches with the suppression of IFN signal seen in panel C. Values reported were normalized to the vehicle control. Different colored markers represent independent experiments.

**C:** Time-course for IFN induction by DTX in BJ fibroblasts. Markers represent the average of technical replicates +/- SEM.

**D:** Minimal cytotoxicity was observed with BAY1217389 after 3 day treatment of BJ fibroblasts measured by ATP amount with Cell-titer Glo assay. Markers represent the average of technical replicates +/- SEM.

**E:** Time-course for IFN induction by BAY1217389 in BJ fibroblasts. Markers represent the average of technical replicates +/- SEM.

**F:** MT stabilizers show similar IFN induction profiles. BJ fibroblasts were treated with Docetaxel (DTX) and Epothilone-B (EpoB), two MT stabilizers with distinct chemical structures. Markers represent the average of technical replicates +/- SEM.

**G:** Dose-responses for MPS1 inhibitors. BJ fibroblasts were treated with MPS1 inhibitors for 3 days and IFN signaling was measured with co-culture assay. BAY1217389, CFI-402257 and AZ3146 showed similar curve shapes with varying EC50 that matches their reported potencies. Interestingly, Reversine shows minimal IFN induction and produces a bell-shaped response, likely due to off-target effects on Aurora Kinase B and cytokinesis interference.

**H:** Cell viability measured after 3 Days of anti-mitotic treatment. MPS1 inhibition reduces cell death from DTX. Markers represent the average of technical replicates +/- SEM.

**I:** The density dependence of IFN induction by MPS1 inhibition and DTX. Optimal density for 4 day assay is between 7-10000 cells per well in a 96-well plate. When BJ fibroblast were seeded too light, or too heavy, the measured IFN signal was reduced. The reduction of IFN signal at higher density is likely due to a reduction in proliferation as cells approach confluency. Markers represent the average of technical replicates +/- SEM.

**J:** The density dependence of IFN induction by 18 hour 2'3'-cGAMP stimulation. Maximal IFN signal was achieved by 50K cells per well in a 96-well plate and did not show a reduction at confluency. Markers represent the average of technical replicates +/- SEM.

**K:** Western blots confirmed STING -/- status of BJ fibroblasts

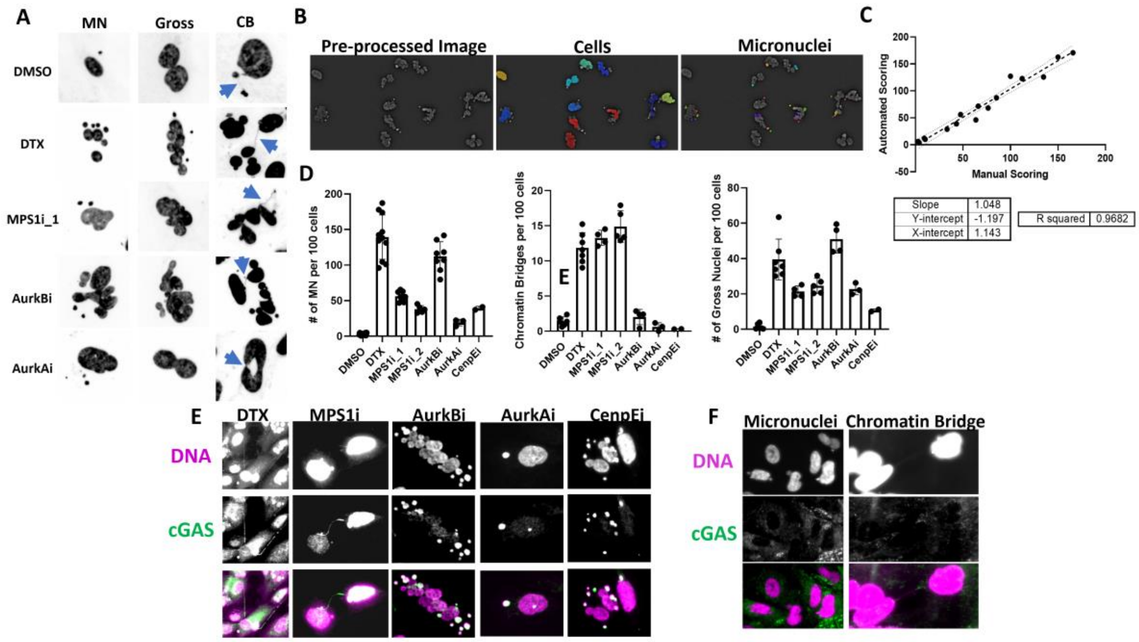


**Supplemental Figure 3: Analysis of Abnormal Chromatin Structures**

**A:** Example nuclear morphologies produced by anti-mitotics. Aberrant structures are classified as micronuclei, gross nuclei and chromatin bridges. Chromatin bridges are highlighted with blue arrows.

**B:** Example automated segmentation results. Images are pre-processed to smooth and enhance edges, they are then fed to the NucleAIzer segmenter and the output is analyzed to differentiate cellular objects from micronuclei.

**C:** Automated segmentation algorithm for # micronuclei/#cell agreed with a manually scored image set.

**D:** Frequencies of different types of nuclear structures produced after anti-mitotic exposure. Each marker represents an independent experiment in which 3-5 FOVs were scored.

**E:** Anti-mitotics produce increase aberrant cGAS-positive chromatin structures. Representative nuclear abnormalities produced by different drugs are shown. Only DTX and MPS1i produce cGAS-coated chromatin bridges.

**F:** cGAS antibody showed no localized signal to abnormal nuclear structures in BJ cGAS KO cells, which confirms the specificity of the antibody.

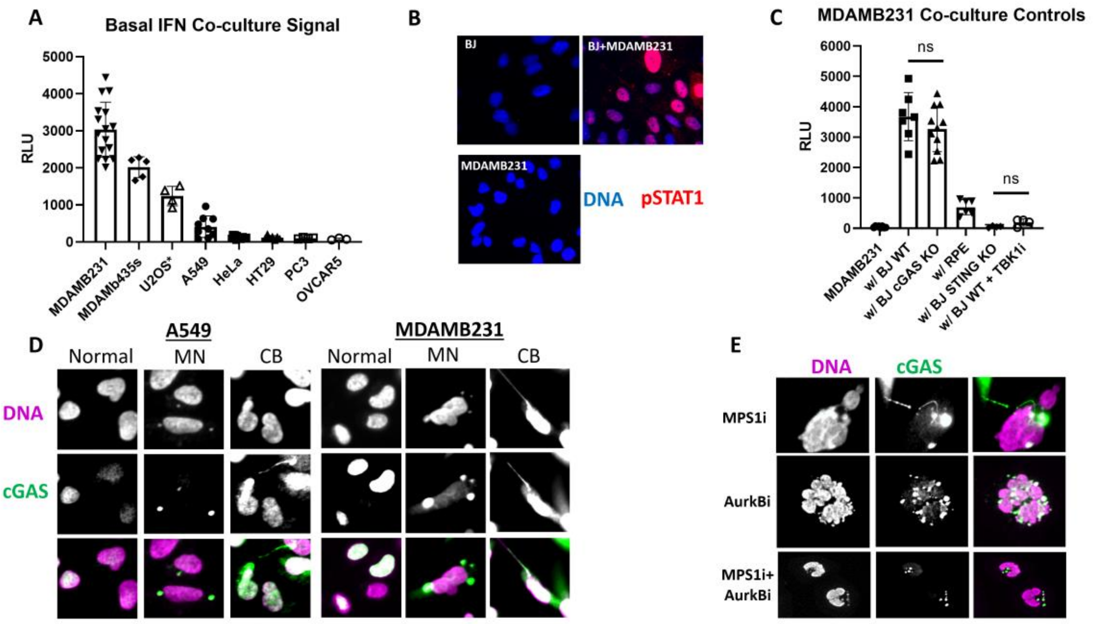


**Supplemental Figure 4: Tri-culture assay to measure cGAS activity in cancer lines**

**A:** IFN detected from co-culturing cancer cell lines with BJ cGAS KO cells. Cells were cultured at a 1:1 ratio with 10K total cells per well. THP1 STING KO reporter cells were added on the third day of co-culture and luminescence was read 18 hours later. Different cancer lines produced different amounts of IFN which indicates various levels of basal cGAS activity. Markers represent independent experiments.

**B:** Co-culturing MDAMB231 cells with BJ fibroblasts drove IFN signaling as assessed by pSTAT1 nuclear translocation.

**C:** Controls for the tri-culture assay. MDAMB231 cells were used to validate the principles of the tri-culture assay. IFN signal from co-culturing MDAMB231 cells with BJ fibroblasts required functional STING in the BJs which implies paracrine cGAMP spread from active cGAS in the MDAMB231 cells. RPE cells, which express very low levels of cGAS but contain functional STING, also produced IFN in response to MDAMB231 cGAMP secretion. Markers represent independent experiments.

**D:** Cancer lines have constitutive levels of atypical nuclear structures which stain positive for cGAS.

**E:** Anti-mitotics induce cGAS-positive micronuclei and chromatin bridges in cancer lines. Only the inflammatory conditions, MPS1i, produced cGAS-positive chromatin bridges.
